# Interplay between Polycomb PCGF protein interactomes revealed by screening under endogenous conditions

**DOI:** 10.1101/2022.05.18.492435

**Authors:** Nayla Munawar, Kieran Wynne, Giorgio Oliviero

**Affiliations:** Department of Chemistry, College of Sciences, United Arabs Emirates University, Al-Ain 15551, United Arab Emirates; Conway Institute of Biomolecular and Biomedical Research, University College Dublin, Dublin 4, Ireland; Systems Biology Ireland, School of Medicine, University College Dublin, Dublin 4, Ireland

## Abstract

The six PCGF proteins (PCGF1-6) define the biochemical identity of Polycomb Repressor Complex 1 (PRC1) subcomplexes. While structural and functional studies of PRC1 subcomplexes have revealed specialized roles in distinct aspects of epigenetic regulation, our understanding of variation in protein interaction networks between the PCGF subunits is incomplete. We carried out an affinity purification mass spectrometry (AP-MS) screen of subunits PCGF1 (NSPC1), PCGF2 (MEL18), and PCGF4 (BMI1), using an immunoprecipitation approach that replicated endogenous cellular conditions in a cell line capable of differentiation programs. Over 200 interactions were found, including 83 that had not been described previously. Bioinformatic analysis found that these interacting proteins covered a range of functional pathways, often focused on cell biology and chromatin regulation. We found evidence of mutual regulation (at mRNA and protein level) between distinct PCGF subunits. Furthermore, we confirmed that disruption of each subunit using shRNA results in reduced proliferation ability. Overall, our work adds to understanding of the role of PCGF proteins within the wider cellular network.

## INTRODUCTION

Chromatin accessibility reflects the degree to which nuclear macromolecules physically compact DNA into a small volume within the nucleus. It is determined by the occupancy and topological organization of nucleosomes as well as other chromatin-binding factors that occlude access to DNA (1). Chromatin-binding factors cooperatively regulate gene expression throughout alteration of chromatin architecture. A class known as chromatin re-modelers can rearrange the accessible or permissive chromatin conformation. These enzymes mainly regulate Post-Translational Modification (PTM) of the N-terminal tail regions of the histone proteins, and include functionally related families of heterooligomeric protein complexes, including the Polycomb Repressive Complexes (PRC) (2) (3) (4). PRC complexes function to regulate gene expression during mammalian development (5) (6) (7).

PRC complexes assemble in two major configurations: Polycomb Repressive Complex 1 (PRC1) is an E3 ubiquitin ligase that mono-ubiquitylates histone H2A at lysine 119 (H2AK119ub1), while Polycomb Repressive Complex 2 (PRC2) houses a methyltransferase that can mono-, di-, and tri-methylate histone H3 at lysine 27 (H3K27me1, H3K27me2, and H3K27me3) (8) (9). Some non-histone substrates (e.g. STAT3, RORα) can also be methylated (10).

The core of the PRC2 enzyme complex is composed of four proteins: EZH1/2, EED, SUZ12, and RBAP46/48 (11) (12). However, accessory PRC subunits have been described, such as AEBP2, JARID2, PCL1 (PHF1), PCL2 (MTF2) and PCL3 (PHF19) (12). Some combinations of these accessory proteins are mutually exclusive, for example JARID2 in combination with most of the other accessory proteins (13) (14).

Biochemical analysis of PRC1 complexes has also revealed a variety of alternative forms. PRC1 complexes consist of a RING1 protein (RING1A or RING1B) and one of six alternative Polycomb Group Ring Finger proteins (PCGF1-6) (8) (15). PCGF1, PCGF2, PCGF3, PCGF4, PCGF5 and PCGF6 are also known as NSPC1, MEL18, RNF3, BMI1, RNF159 and MBLR respectively. PRC1 complexes can be classified into canonical or non-canonical forms (cPRC1 and ncPRC1) (8). Both complexes mediate histone H2A monoubiquitination via the E3 ubiquitin ligase component RING1A/B. In addition, cPRC1 complexes contain CBX (chromobox) proteins that target the complex trimethylated lysine 27 on histone 3 (H3K27me3).

In ncPRC1 complexes, RYBP or YAF2 recognize the H2AK119ub1, resulting in a form of positive feedback (16). In general, cPRC1 complexes are associated with chromatin condensation events, while ncPRC1 are linked to stronger ubiquitination activity (8). In mouse embryonic stem cells, PRC1-dependent H2AK119ub1 leads to recruitment of PRC2 and H3K27me3 to effectively initiate a polycomb domain. This activity is relative restricted to the ncPRC1 variant PCGF1-PRC1 complex that recognizes non-methylated DNA in CGIs by the CxxC-ZF domain of KDM2B (17). This contributes to histone H2A lysine 119 ubiquitylation and gene repression (17) (18) (19).

Polycomb proteins coordinate major differentiation and developmental proteins in many cellular contexts. PRC components have been identified as positive regulators of mESC self-renewal and differentiation pathways (7) (20) (21). Additionally, abnormal PRC protein expression and/or mutation can lead to impaired signaling that inhibits tumor suppressor activity, or promotes proto-oncogene activity, leading to loss of cell identity (9) (22) (23). Hence, much research related to Polycomb-mediated gene repression has employed mouse embryonic stem cell (mESC), a powerful yet accessible model (24) (25) (20).

Pluripotent embryonic carcinoma cell lines, such as NTera-2/cloneD1 (NT2), are an important tool for studying pluripotent and stem cell-like differentiation programs in a human model (26) (27) (28). Upon treatment with retinoic acid (RA), NT2 cells can be induced to differentiate into neuron-like cells, which display a variety of neurotransmitter phenotypes (29) (30) (31).

We previously investigated the PRC1 complex in the NT2 cell model. We focused on the role of PCGF1/NSPC1, a subunit of ncPRC1 complex that functions to maintain the embryonic cell fate by interacting with pluripotency markers such as OCT4, NANOG and DPPA4 (32). The combination of affinity purification and high resolution/high mass accuracy mass spectrometry allows the mapping of protein interaction networks in unprecedented detail (33) (34).

We extended this investigation to include the PCGF proteins PCGF2 and PCGF4. While these proteins share significant amino acid homologies, they form distinct interaction networks in NT2 cells. Notably the two cPRC1 proteins, PCGF2 and PCGF4, each interact with a subset of unique proteins in addition to a shared set, despite sharing 64% amino acid sequence homology. We report that despite all three PCGFs sharing a certain degree of protein homology, this does not imply that they share similar biological functions.

## RESULTS

### A physical interaction screen for PRC1 components purified under endogenous conditions

We used an immunoprecipitation approach, combined with high-resolution mass spectrometry, to identify physical protein interactions of PRC1 components in NT2 cells, while avoiding artefacts arising from overexpression (Figure 1A). Briefly, nuclear lysates were immunoprecipitated with anti-PCGF1, anti-PCGF2, anti-PCGF4, with anti-Rabbit IgG antibody used as negative control. We also included a screen of anti-RNF2/RING1B, the catalytic enzymatic core of PRC1 complexes. This protein is therefore an interactor with all PRC1 complexes containing PCGF subunits. Each lysate was subsequently digested using trypsin immobilized on agarose beads to yield soluble peptides. The peptides were desalted, adsorbed onto C18 zip tips, eluted in high acetonitrile, and separated online by nano-chromatography interfaced with a Q Exactive mass spectrometer (Supplementary material). Lastly, mass spectrometry raw data were used to identify and determine the relative abundance of the proteins, using the MaxQuant platform (35).

**Figure 1.**
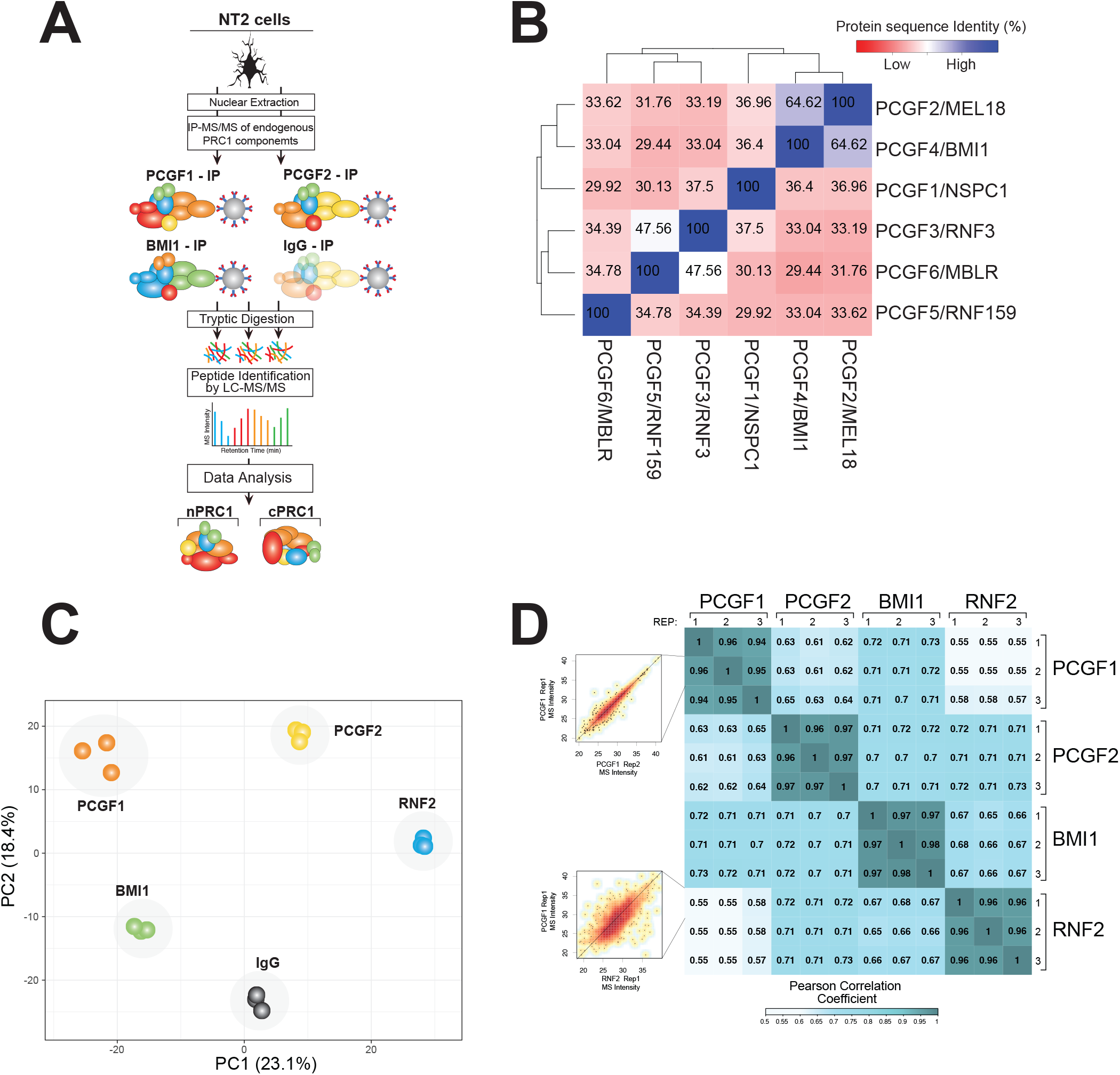
A physical interaction screen for PRC1 components. A: Four subunits of PRC1 subcomplexes were purified under endogenous conditions using immunoprecipitation (IP). Physically interacting proteins were identified and quantified using Orbitrap mass spectrometry. B: Amino acid homology between all six human PCGF proteins were compared using Clustal-Omega. C: Sensitivity and reproducibility of biological replicate and individual IP experiments were compared by plotting a matrix of pairwise Pearson correlation coefficients. D: The physical interactomes of each IP experiment (including biological—replicates) were compared using Principal Component Analysis.

Two protein domains are shared across all the PCGF subunits: RING finger and RAWUL domains (36). These domains likely play an important role during assembly of the PRC1 complexes (37) (38). Comparison of the amino acid sequences of the PCFG variants using multiple alignment (39) shows that the highest sequence homology among them is between PCGF2 and PCGF4 (Figure 1B). Interestingly, PCGF2 and PCGF4 also contain extended C-terminal regions that may be associated with disorder (40) (41). Weaker levels of homology were observed between other PCGF components.

We carried out preliminary experiments to confirm that our approach was sensitive and reproducible. Comparison of mass spectrometry peptide intensity signals for replicate (Biological) repeats confirms that experimental reproducibility was high. The average Pearson correlation between replicates was ~0.95 for biological replicates (Figure 1C). The replicate experiments showed a high degree of correlation. Between immunoprecipitations, reasonable correlation (~ 0.6 - 0.7) observed for the three PGCF proteins, while the single non-PCGF protein analysis (RNF2) showed slightly lower correlation (~ 0.5), as expected.

In order to evaluate the overall dataset, we used principal component analysis (PCA) using the summed peptide mass spectrometry signal intensities for each identified protein as input (‘Label Free Quantitation’ value from MaxQuant) (42) (Figure 1D). Reassuringly, each immunoprecipitation experiment was well separated while the biological experimental replicates were located close together.

### AP-MS screening reveals common and distinct interactomes among PRC1 component proteins

To assess the distinctive PCGF subunit interactomes, we compared protein abundance in samples immunoprecipitated using α-PCGF1, α-PCGF2, α-PCGF4, and α-RNF2 (RING1B) to samples immunoprecipitated using IgG as a negative control (supplementary material). The specificity and effectiveness of the antibodies was confirmed using co-immunoprecipitation experiments (Figure 2A and supplementary material). Each antibody immunocaptured it’s cognate bait protein, while all three baits co-precipitated the expected common PRC1 complex subunit RING1A and RING1B an E3 ubiquitin ligase for H2AK119ub and essential component of PRC1 in mammals (43) (44).

**Figure 2.**
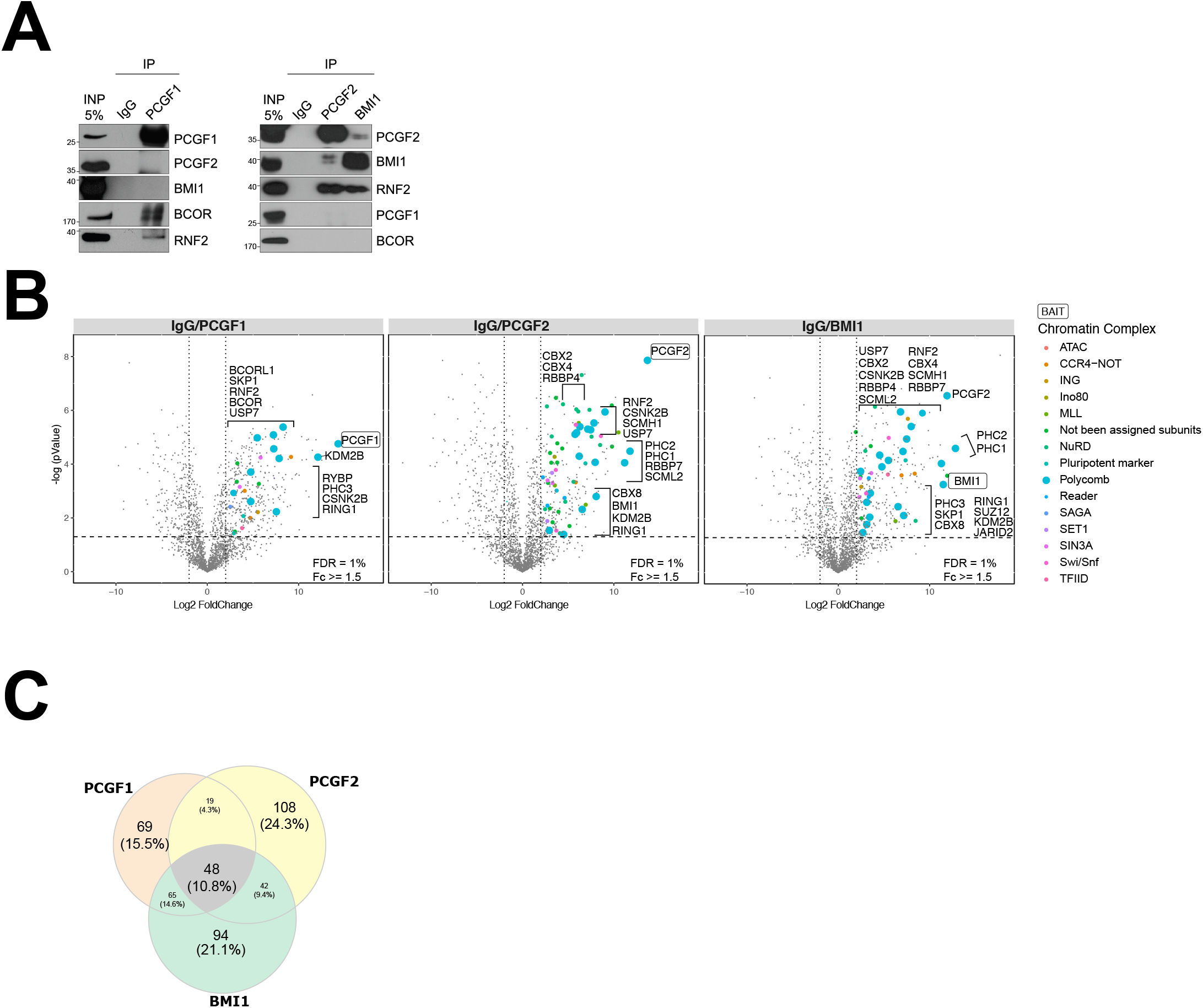
AP-MS screening reveals common and distinct interactomes among PRC1 component proteins. A: The specificity of the antibodies used in the PCGF1, PCGF2, and PCGF4 immunoprecipitations were confirmed using western blotting. Positive (the cognate proteins, and the PRC1 core subunit RING1A/RNF2) and isoform-specific proteins (BCOR) behaved as expected. Some overlap in specificity between PCGF2 and PCGF4 was observed. B: The set of identified proteins for each immunoprecipitation experiment was projected onto volcano plots to identify statistically robust hits. C: Venn diagram showing the range of overlap among the physical interactomes of PCGF1, PCGF2, and PCGF4, as well as subsets uniquely identified for each bait.

To support our strategy, we investigated the RNF2 interactome as PRC1 quality control. In line with previous PRC1 interactome analyses (45) (46), RNF2 was immunoprecipitated with all six PCGF subunits (supplementary material). Volcano plots were used to project each protein onto a chart showing enrichment in each immunoprecipitation experiment relative to the IgG control (x-axis) versus the significance of that finding based on the t-test (y-axis) (Figure 2B).

A total of 48 interactions were shared across all PCGF proteins (Figure 2C). PCGF2 and PCGF4 shared 42 interactions, while PCGF1 and PCGF2 shared 19 candidate interactors, and PCGF1 and PCGF4 had 65 interactors in common. In general, the proteins observed to interact in NT2 cells correlated well with those reported by Hauri, Gao and Wiederschain *et al.* (45) (46) (47), suggesting that the composition of PRC1 complexes are reasonably conserved despite different cellular contexts (Supplementary material).

The BCOR component of the non-canonical PRC1 complexes was precipitated as expected by anti-PCGF1, while the canonical complexes were not. The PCGF1 interactome also exhibited non-canonical PRC1 co-factors, such as BCORL1 and SKP1.

The PCGF2 and PCGF4 interactomes contained the chromo-domain protein CX2, CBX4 and CBX8, as expected for canonical PRC1 complexes.

As with all classification attempts, these PCGF-PRC1 variants need to be simplified somewhat. One can gain an insight into the difficulties encountered in correctly categorizing these complexes upon consideration of the number of cell types in conjunction with the multiple protein characterization strategies employed. It is also important to mention that PRC1 complexes are highly dynamic structures that evolve in tandem with progression between cell states (46) (48). In this study, we aimed to elucidate the PCGF-PRC1 architecture in the presence of auxiliary subunits, classified as either non-canonical or canonical PRC1 employed affinity proteins purified in native conditions.

In line with a certain degree of protein homology divergence across all PCGF subunits, we reported that the BCOR component of non-canonical PRC1 complexes is precipitated as expected by anti-PCGF1. We subsequently observed that the PCGF2 and PCGF4 subunits co-purified and shared the same chromo-domain proteins and ubiquitin ligase modules, as expected for canonical PRC1 complexes. Overall, we observed that the PCGF interactomes contained a heterogeneous collection of subunits, sometimes in a sub-stoichiometric manner, demonstrating the differences inherent in the PRC1 architecture. In addition, the native PCGF interactome had not completely elucidated.

We compared our data with the Gao and Hauri *et al* studies (45) (46), using a Venn diagram to determine common and unique PRC1 features obtained from diverse protein characterization strategies (supplementary material). We found that all the PCGF-candidate interactors reported by Gao and Hauri *et al* were also identified by us, being shared across the three studies. We also observed interaction candidates not yet described: 191 in the PCGF1, 207 in the PCGF2, and 237 in the PCGF4 interactome (Supplementary material).

The differences between the two studies may be due to the analytical strategies performed and/or the context of the biological samples being analyzed. In contrast to our use of NT2 cells, Gao *et al* used an affinity-tagged strategy in 293TREx cells (45). A similar strategy was employed by Hauri *et al* (46) who generated stable HEK293 cell lines exhibiting a tetracycline-inducible expression of several polycomb group components.

### Stoichiometry and molecular mass of the isolated PRC1 complexes

Many of the interactions we identify are components of other chromatin re-modelers (Figure 2B and supplementary material).

By retrieve the most recent depository chromatin remodelling complex (49) (50) (51) we specifically detected the following: Ada2a-containing (ATAC) ATAC; Carbon catabolite repression (CCR4) negative; inhibitor of growth (ING); mixed lineage leukemia (MLL); nucleosome remodeling and deacetylases (NuRD); Spt-Ada-Gcn5 acetyltransferase (SAGA); SET domain-containing protein (SET); histone deacetylase complex subunit (SIN3A); SWItch/Sucrose Non-Fermentable (SWI/SNF); and the general transcription factor IID (TFIID). These results are in line with Hauri *et al*., who reported PRC1 and PRC2 co-purified with several chromatin remodeling subunits encompassing MLL, NSL, ADA2/GCN5/ADA3 transcription activator, NURF, NURD, and SIN3 complexes (46).

We also distinguished unique and different chromatin re-modelers which are not yet assigned, such as a CCR4-NOT complex uniquely co-purified with PCGF1. We also observed CNOT1 and CNOT4 subunits. Both CNOT1 and CNOT4 are involved in E3 ligase activity and promote histone ubiquitination (52) (53). In addition, we identified other subunits such as ARID2, ATRX, BRD7, SMARCB1, SMARCA4, SMARCC1, SMARCC2, SMARCD1, SMARCD2, and SMARCE1 (54) (55). The mechanism by which BAF complex disengagement leads to polycomb repressor complex-driven re-establishment of heterochromatin signatures associated with gene repression is still not defined (55). We did not perform any further immunoblotting validation for these novel candidates due to the lack of highly specific antibodies against them.

Since PRC1 is itself a high molecular weight (MW) multiprotein assembly, and these potential interactors are themselves components of multiprotein assemblies, we next focused on analyzing the physical form of the PRC1 complexes that we isolated. First, we estimated stoichiometry by dividing the normalized mass spectrometry intensity signal for each protein (LFQ) by the protein MW (iBAQ score) (56) (Figure 3A). Core members of the PRC1 complex (such as RNF2, RING1A, KDM2B, USP7, CSNK2B, and PHC3) that are shared among all three PCGF-PRC1 interactome complexes were present in approximately equal stoichiometry. PCGF2-PRC1 and PCGF4-PRC1 variant interactomes exhibited a common stoichiometric pattern profile and shared canonical PRC1 subunits including the following chromobox proteins: PHC1, PHC2, CB2, CBX4, CBX8, SCML2, SCMH1, RBBP4, and RBBP7.

**Figure 3.**
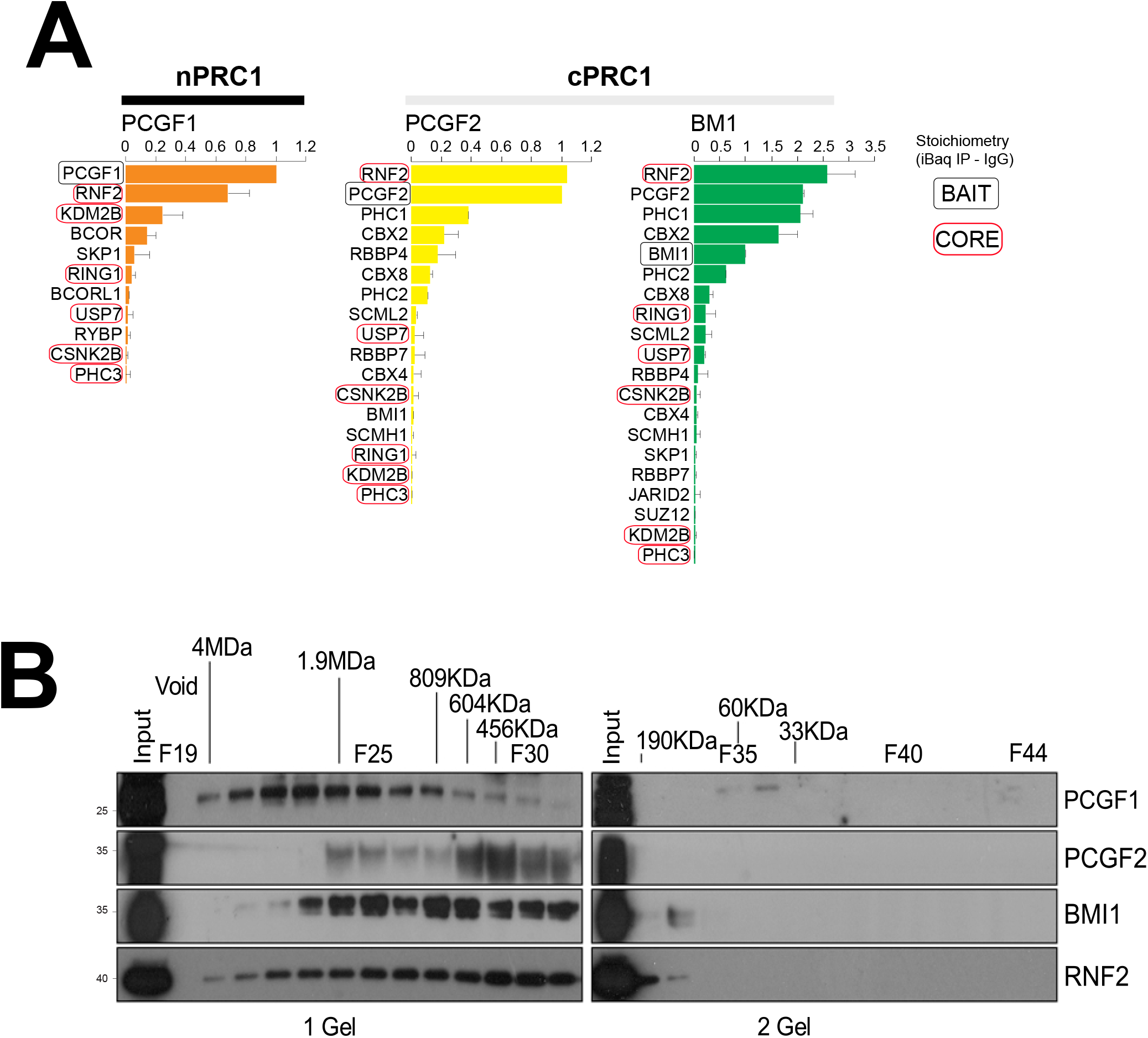
Stoichiometry and molecular mass of the isolated PRC1 complexes. A: The stoichiometric contributions of individual subunits to each IP was estimated using peptide signal intensity adjusted for protein size. Relative quantities were normalized to the signal recorded for bait peptides. B: Nuclear lysates separated using gel filtration were run as fractions on SDS-PAGE and probed using antibodies to PCGF1, PCGF2, PCGF4, and RING1A/RNF2.

To investigate the molecular mass of the isolated complexes, nuclear protein lysate from NT2 cells was separated by size exclusion chromatography and the fractions probed using antibodies against PCGF1, PCGF2, PCGF4, and RNF2 (Figure 3B). All four PRC1 component proteins were found to be present in high mass complexes. These varied greatly in size, from 200KDa to 4MDa. The size exclusion experiments confirms that the high mass complexes that contain PCFG2 and PCGF4 are largely overlapping, while the PCGF1 profile seems to belong to a higher mass range complex.

### Functional enrichment of PCGF interactomes map to multiple pathways

Next, we asked if the observed interactomes were associated with particular molecular pathways. We used the Gene Ontology (GO) analysis (“BP”, “biological process”) (Figure 4A) to investigate this. We first analyzed the whole PCGF interactome to create a comprehensive overview mapping of pathways associated with the PCGF interactomes. The functional categories are displayed in a dot plot cluster and represent the significant biological process enriched. Overall, five main categories were found to be significantly enriched: “mRNA splicing”, “regulation of chromosome organization”, “histone ubiquitination”, “regulation of G0 to G1 transition”, and “histone monoubiquitination”.

**Figure 4.**
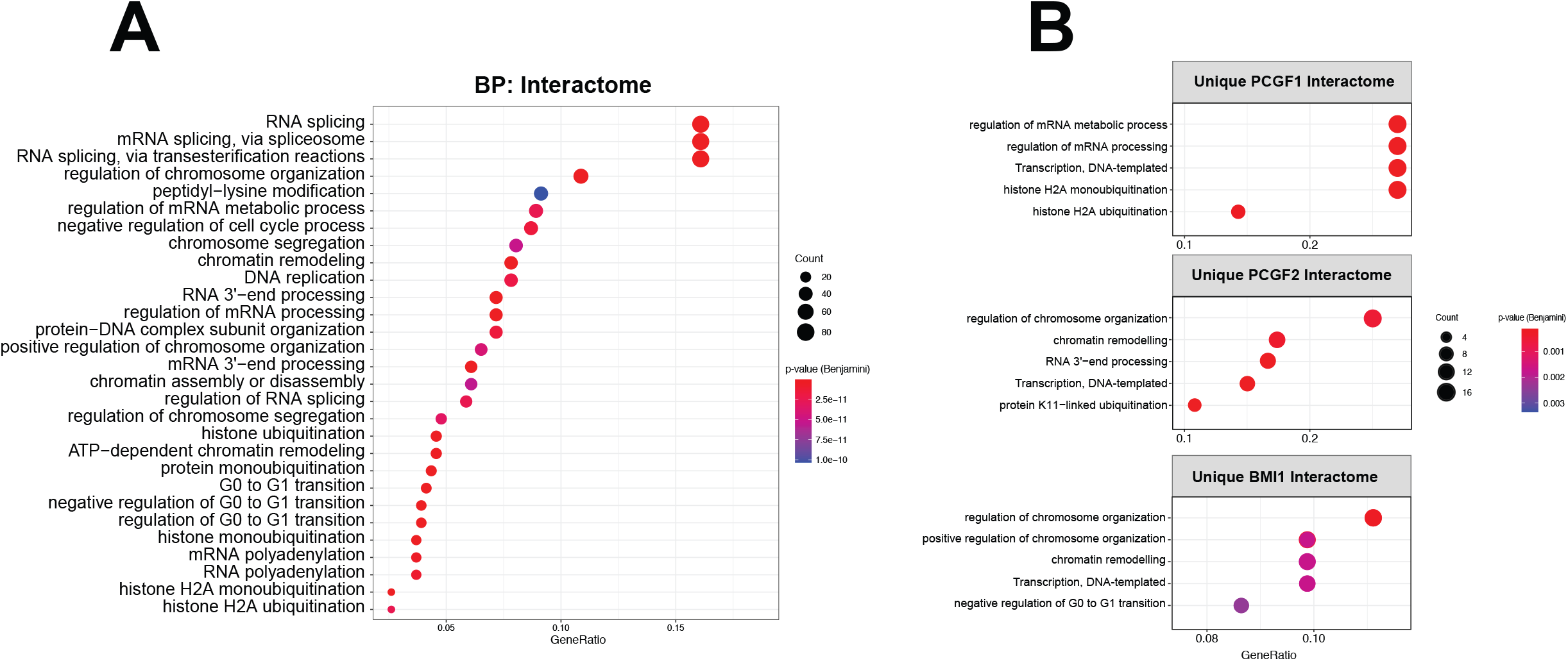
Functional enrichment of PCGF interactomes map to multiple pathways. Gene Ontology (GO) analysis was used to identify enriched pathway components in the overall (A) and individual PCGF interactomes (B). Categories are sorted by p-value (Spearman’s Rank Correlation Coefficient), while the dot size represents the number of proteins correspondent to the source pathway.

We performed similar analysis among the distinctive PCGF interactomes to assess the unique pathways which may affect the PRC1 organization through their unique PCGFs features (Figure 4B). The PCGF2 and PCFG4 interactomes exhibited similar biological functional properties, including common biological annotations such as “regulation of chromosome organization”, “transcription, DNA-templated”, and “chromatin remodeling”. We also observed cell cycle-related terms such as “negative regulation of G0 to G1 transition” as uniquely enriched pathways in PCGF4 interactomes, while “histone ubiquitination” categories were shared between the PCGF1 and PCGF2 interactomes. Overall, we observed distinctive PCGF interactomes which revealed a different biological function related to each PCGF interactome. We also confirmed associated functional biology concepts that are known to be Polycomb-related.

### The role of PCGF subunits in NT2 cells

In order to assess the functional effect of disrupting PCGF expression in NT2 cells, we carried out a knockdown screen for the PCGF subunits. Successful knockdown of each PCGF variant was achieved using the shRNA method (Figure 5A). As expected, global levels of H2BK119ub were reduced, most prominently for PCGF4, in line with previous reports (57). Furthermore, levels of PCGF4 mRNA itself were reduced following knockdown of PCGF2, and vice versa. This suggests that PCGF2 and PCGF4 may influence their regulation both at transcriptional and protein levels. This corroborates previous evidence of a synergistic requirement for these PcG proteins in the maintenance *Hox* gene expression during early mouse development (58). Mouse embryos deficient for PCGF2 and PCGF4 exhibit similar posterior transformations of the axial skeleton and display severe immune deficiency (58).

**Figure 5.**
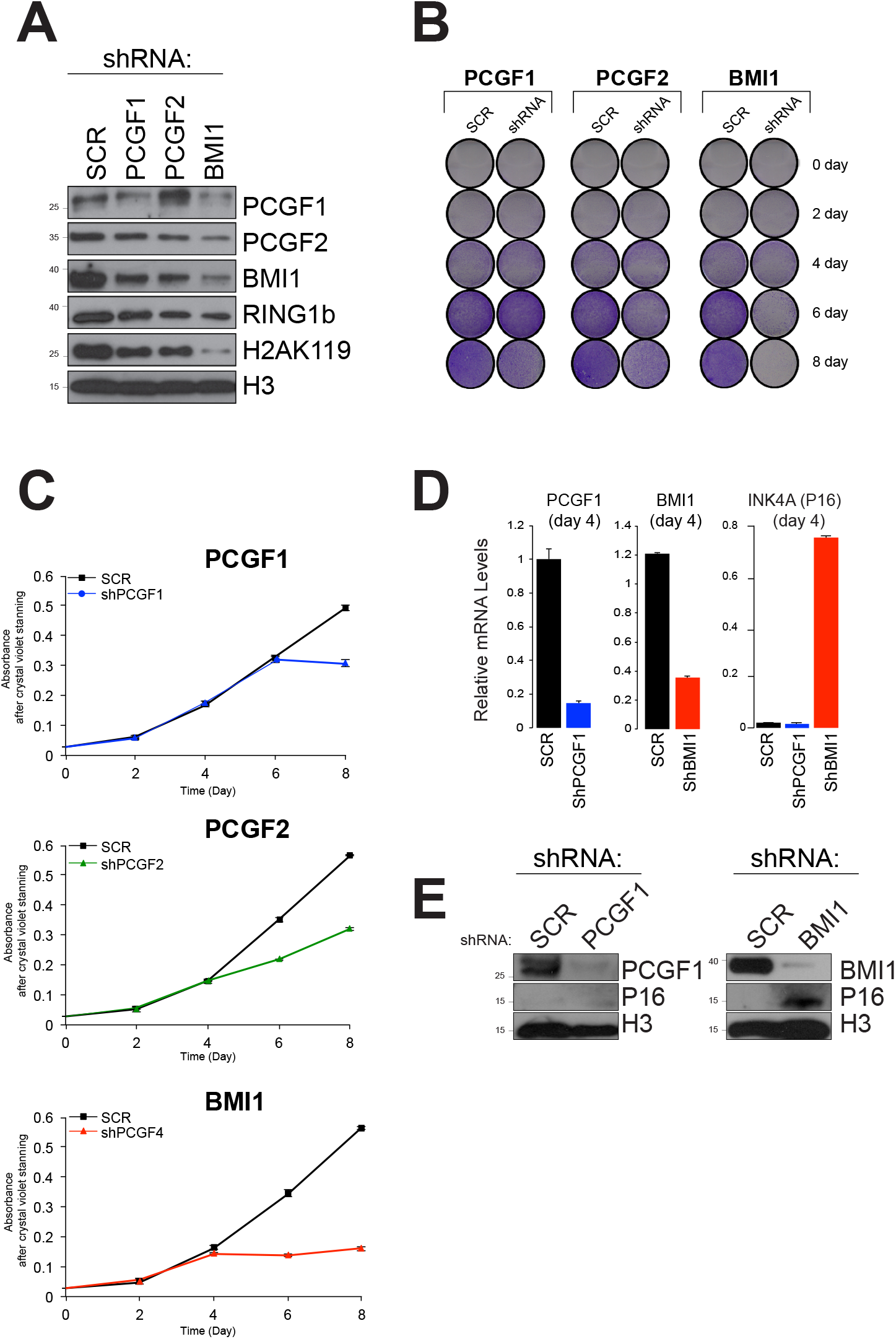
Functional enrichment of PCGF interactomes map to multiple pathways. A: Western blotting was used to confirm successful partial knockdown of protein expression for PCGF1, PCGF2, and PCGF4. Ubiquitination of histone H2A K119 was also reduced. B: Cell proliferation was quantified using the crystal violet assay. C: This assay allowed comparison of the kinetics for each knockdown, with PCGF4 showing a more pronounced reduction of proliferation rate than PCGF1 or PCGF2. D: The INK4A-P16 marker of cellular senescence was selectively reduced by treatment with PCGF4 but not PCGF1 shRNA. E: This latter result was confirmed by western blot.

Conversely, some minor effects of PCGF1 protein expression alteration seem to be detected during PCGF2 downregulation. These results suggest an auto-regulatory activity among many, or even all, PCGF genes. In order to observe differences at a phenotype level during the depletion of PCGF subunits, we performed cell viability experiments over a time course using the crystal violet assay (Figure 5B). We monitored the cell growth rate in the presence or absence of PCGF gene expression. We compared and screened the most efficient shRNA construct on HEK293 cells and selected those shown to have a higher knockdown efficiency to the corresponding PCGF subunits (supplementary material material).

We subsequently generated PCGF lentivirus for quantitative assessment of gene knockdown expression in NT2 cells (Figure 5C). We observed that cell growth rate is reduced after four days following disruption of PCGF4 expression. A similar trend, although slightly less pronounced, was observed for PCGF2 depletion, while in PCGF1 samples the cell proliferation rate did not change (Figure 5 B,C). The observation that PCGF4 can influence growth rate raises the possibility of a mechanism related to senescence. In line with a previous study, regulatory mechanisms yielded by PCGF4 expression controls the cell cycle through the regulation of the *Ink4a/Arf* locus (59) (60). To confirm this, we measured the mRNA and protein levels of *p16*, a senescence marker (61) (62), at protein and transcriptome level respectively (Figure 5E). Interestingly, only PCGF4 was found to influence the cell viability through *p16* expression (Figure 5 D,E). This may suggest that other PCGF proteins can influence cell growth via independent, non-senescent pathways.

## DISCUSSION

In the last decade the PRC1 complex has been intensively investigated (63) (64). A recent study involved the dissection of the PRC1 assembly composition and indicated that six different PRC1 variants existed. Each PRC1 variant exhibited a different PCGF subunit, indicating that PRC1 complexes contain mutually exclusive homologs of the PCGF protein (15) (45) (65).

There are several mammalian PRC1 complexes, each characterized by a single homolog (in different combinations) of the *Drosophila* proteins Psc (PCGF1-6), Ph (PHC1-3), Pc (CBX2, 4, 6, 7, and 8), Sce (RING1A/RNF2), and Yaf2 (RYBP). Canonical PRC1 complexes also contain homologs of the chromatin reader protein, CBX, which exhibited chromo-domains and are recruited to chromatin via their ability to bind the PRC2-mediated H3K27me3 mark (8) (66) (67).

Non-canonical PRC1 complexes, which do not contain CBX subunits, are able to recognize the Polycomb target gene through other auxiliary proteins, such as BCOR or KDM2B (18) (68) (69). However, all PRC1 complexes contain the RNF2/RING1b and RING1/RING1a E3 ubiquitin ligase enzymes that target histone H2A lysine 119 for mono-ubiquitination (H2AK119ub) (68) (69). This histone mark co-localizes genome-wide with the PRC2-mediated H3K27me3 histone mark and is associated with chromatin compaction and promotion of gene silence (64) (70).

We investigated the physical interactomes of selected PCGF subunits. PCGF1, PCGF2, and PCGF4 showed a degree of protein sequence homology and were expressed in NT2 cells. We didn’t attempt to investigate the other PCGF proteins due to the lack of commercially available antibodies. In NT2 cells, the protein dynamic range analysis suggested a lower protein expression level for PCGF3, PCGF5, and PCGF6. Accordingly, the PCGF-PRC1 architecture may not reflect the native organization (supplementary material).

Previous studies identified two domains shared across all the PCGF proteins: RING finger and RAWUL (37) (36). The contribution made by the RING finger and RAWUL domains may play an important role in defining PRC1 composition. How non-canonical PCR1 subcomplexes are recruited to chromatin remains less understood (63). In B-cell lymphoma 6 (BCL6), the interacting co-repressor (BCOR) forms a complex with RING1/RNF2, RYBP, PCGF1, and KDM2B, causing transcriptional repression during lymphocyte development (69) (71). The direct binding partner of PCGF1 is BCOR, which has emerged as an important player in development and health (38) (72) (73). Recruitment of non-canonical PRC1 complex to chromatin depends on its KDM2B subunit, which can recognize unmethylated CpG islands (74) (68). The activities of non-canonical PRC1 complexes showed that recruitment of the PCGF1-PRC1 variant results in H2AK119 ubiquitylation. This can promote the recruitment and/or stabilization of PRC2 to the chromatin and reinforce the deposition of H3K27me3 (18).

In mammals and in *Drosophila*, PCGF2 and PCGF4 share most of their Polycomb subunits including a group of related proteins, termed Polyhomeotic (PHC1, 2, and 3) (75). PCGF2 and PCGF4 are considered to form canonical-PRC1 complexes and bind to chromatin via CBX proteins that recognizes H3K27me3 (8) (67).

Recently, the role of PRC1-6 subcomplexes was studied by combining the development of highly specific PCGF1-6 antibodies in-house, with the generation KO mESC lines depleted for all six PCGF proteins (15) (65). The genome-wide occupancy of all PRC1 subcomplexes was mapped to determine their functional control in pluripotent cell modelling. The results suggested that the activities of PCGF1 and PCGF2 are strongly linked with transcriptional repression and display extensive functional overlap (15). In mouse cells, the PCGF2 and PCGF4 double mutant embryos exhibit severe growth retardation, accelerated apoptosis, and defects in the maintenance of stable gene expression during early development.(58)

PCGF2, is also involved in cell proliferation, differentiation, and embryogenesis (41). PCGF2 is a target of the protein kinase AKT (76). AKT phosphorylates PCGF2 to disrupt the interaction between PCGF2 and other PRC1 members that cause tumorigenesis in breast cancer (60) (77). Missense substitutions of the Pro65 residue of PCGF2 shows severe clinical outcomes recognizable as developmental delay, intellectual disability, impaired growth and several brain, cardiovascular, and skeletal abnormalities (41). PCGF2 has also been found to be essential for ESC differentiation into early cardiac-mesoderm precursors, and exhibits a distinctive PCGF2-PRC1 activity to control the expression of the negative regulators of the BMP pathway and genes involved in cardiac development (48).

PCGF4 was the first PCGF protein identified based on its ability to act as an oncogene, collaborating with c-MYC in a transgenic model of a mouse lymphoma (78) (79). Further studies have attributed this oncogenicity to the ability of PCGF4 to directly repress the *INK4A/ARF* gene locus, which encodes the tumor suppressors p16INK4A and p14ARF (59) (60). The repression of these two genes leads to increased proliferative capacity and delayed senescence in mammalian cells. Bmi1-null mice display severely impaired stem cell self-renewal in the neural, mammary, and hematopoietic lineages (58) (80) (81).

These phenotypes are largely associated with a failure to repress the expression of the *Ink4a/Arf* locus, although co-deletion of this locus leads to only a partial rescue of the stem cell self-renewal phenotype (81). Furthermore, in a mouse model of glioma, Bmi1 expression was shown to enhance tumor progression even in *Ink4A/Arf*-null cells (82). This data suggests that BMI1 has functions in cancer development and stem cell biology, independently of the *p16INK4A* and *p14/p19ARF* pathways. Furthermore, PCGF4 is required for the self-renewal of NSCs in the peripheral and central nervous systems, but not for their proliferation or differentiation (83).

In our study we evaluated the role of PCGF-PRC1 organization in NT2 cells, a cell therapy model for the investigation of human neurogenesis, regeneration, and drug screening (26) (27) (28). For over two decades, the role of Polycomb-mediated gene repression has been dissected mostly in mouse embryonic stem cells (mESC), which is considered a “gold standard” model for epigenetics research (24) (25) (20). The focus on this method has limited Polycomb characterization in other cell models, such as cell types that reflect the tumorigenesis environment or exhibited cancer genotype and heterogeneity. We asked if differences in PCGF-PCR1 composition may be reflected by divergence at the protein sequence level, or if they may be influenced by differences at the protein-protein interaction level.

We favored an endogenous immunoprecipitation approach of analyzing PCGF1, PCGF2, and PCGF4 subunits to characterize the behavior of the PCGF-PRC1 variant in a manner as close to the native cell condition as possible, and particularly to avoid artefacts arising from exogenous expression affinity-tagged form. Subsequently, we applied a label-free mass spectrometry strategy to dissect the PCGF-PRC1 variant assembly and distinguish individual interactors that associated preferentially in only one PCGF-PRC1 variant, in two PCGF-PRC1 variants, or in all three PCGF-PRC1 variants.

We observed a significant degree of common interactions among each purified PCGF, consistent with previous reports by Gao, Hauri, and Wiederschain *et al.* This suggests that the composition of PRC1 complexes is reasonably conserved, despite a difference in cellular contexts. Our study also found novel interactors. We favored an endogenous immunoprecipitation approach to mimic the native physiological cell environment. In contrast, previous studies have employed the tandem affinity purification method, which may partially explain the differences between the two strategies (84).

Ectopic protein expression following tandem affinity purification was unable to isolate and identify interacting proteins in 22% of purified tagged proteins in yeast proteome (85). The intrinsic quality of the TAP tag may affect the affinity binding efficiency. Therefore, a relatively low efficiency of purification can be observed. This may explain the large amount of novel chromatin remodeling subunits detected in our strategy, in comparison with previous reports.

The TAP tag added to a target protein may interfere with protein function, location, and complex formation, which is particularly relevant between PCGF2 and PCGF4 as they share sequence homology. Another limit of the TAP strategy comes from the competition of endogenous proteins with the tagged protein, especially if the tagged protein is located in a protein complex, which may explain why PCGF2 and PCGF4 were not present in a complex with each other.

PcG was originally described as a set of genes responsible for controlling proper body segmentation in *Drosophila* (86). Subsequently, the function of PcG was dissected in mammalian models and shown to play a crucial role in regulation in stem cells and embryonic development (7). This supports the selection of NT2 cells as models to simulate stem cell characteristics, which may explain the detection of several chromatin re-modelling subunits mainly identified as master regulators of gene expression via chromatin modifications and compaction.

We then dissected the distinctive functional biological of the PCGF-PRC1 assembly through Gene Ontology (GO) analysis, to create a comprehensive map of the chromatin pathway environment and to improve gene-annotation enrichment analyses related to chromatin environment (which was poorly annotated). We reported cell cycle-related terms such as “negative regulation of G0 to G1 transition” as uniquely enriched pathways in the PCGF4 interactome, while, as expected, “histone H2A ubiquitination” related categories were shared across all three PCGF4.

Lastly, we investigated the role of PCGF subunits in gene regulation of NT2 cells using knockdown screening against the cognate PCGF subunits. We observed through cell viability assay that PCGF4 uniquely affects cell proliferation, rather than the other PCGF auxiliaries.

Despite PCGF2 and PCGF4 sharing amino acid sequence homology and overlapping interactomes, only PCGF4 directly regulated the expression of *INK4a/ARF.* Possible support for this idea was observed in Morey *et. al.* (48), where expression of PCGF2 gradually diminished upon differentiation in mESCs, while in contrast the protein level of PCGF4 upregulated. Pluripotent mESC does not express detectable levels of PCGF4 in either protein or transcriptional levels, and forced PCGF4 expression had no obvious influence on mESC self-renewal (87).

Overall, our experiments yielded important insights into the composition of PCGF-PRC1 assembly complexes and linked alternative PRC1-related complexes to distinct molecular functions. Importantly, we showed a unique link between PCGF4 and p16 expression, potentially linking this protein (and hence the PCGF4-PRC1 complex) to the process of senescence.

In line with previous studies, unique complex components have also been identified for different PCGF homologs, which suggests that they are not completely redundant and that they may also have some independent functions (15) (45). Further insights into the genome-wide localization and complex composition of variant PRC1 complexes in different cellular contexts will likely add to our understanding of their individual and overlapping functions and contribute to our understanding of the PRC1 assembly.

To date, the notion that sequence homology implies functional similarity through common interactor partners is still not proven. Furthermore, how protein similarity may influence the organization of a chromatin re-modeler has still not been elucidated. The investigation of the proteome interactome landscape appears to be a very important distinguishing factor in the definition of the degree of protein similarity in protein families. The integration of protein homology analysis and affinity purification, followed by mass spectrometry analysis, may open a new avenue in solving one of the central problems in modern biology: the aim of identifying the complete set of protein interactions in, and important biological processes of, a cell, including catalyzing metabolic reactions, DNA replication, DNA transcription, responding to stimuli, and transporting molecules from one location to another.

## MATERIAL AND METHODS

### Cell culture

NTera-2/cloneD1 (NT2) cells (ATCC, CRL-1973) were cultured in 92mm tissue culture dishes Nunclon (Fisher Scientific) in Dulbecco’s Modified Eagle Medium (DMEM) supplemented with 10% (v/v) Fetal Bovine Serum (Hyclone), 100U/ml penicillin and 100U/ml streptomycin (Gibco). Cells were passaged by trypsinizing with 0.25% Trypsin-EDTA (Invitrogen) and plated at a ratio of 1:6. For Lentivirus generation HEK293T cells were grown in DMEM medium supplemented with 10% (v/v) FBS (Hyclone), 100U/ml-1 penicillin, and 100U/ml-1 streptomycin (Gibco) and plated at a ratio of 1:10.

### Isolation of Nuclei

Harvested NT2 cells were washed in PBS and resuspended in Lysis buffer (25mm Tris·HCl pH 7.6, 150mm NaCl, 1% Nonidet P-40, 1% sodium deoxycholate, 0.1% SDS, 2μg/ml Aprotinin, 1μg/ml, Leupeptin, 10mm PMSF). The lysates were incubated for 15 minutes on ice and cell membranes disrupted mechanically by syringing 5 times with 21G narrow gauge needle and sonicating at 3Å~ for 2 seconds at high power. Lysates were incubated on ice for another 15 minutes and cleared by centrifugation at 20,000 (RCF) at 4°C for 30 minutes. To harvest the nuclear fraction, lysates were resuspended in an equal volume of Nuclear Buffer (20mm HEPES pH 7.9, 0.2mm EDTA, 1.5mm MgCl2, 20% glycerol, 420mm NaCl, 2μg/ml Aprotinin, 1μg/ml Leupeptin, 10mm PMSF) and dounced 20 times with tight pestle type B. Lysates were incubated for 45 minutes rotating to dissociated chromatin-bound proteins and precleared by centrifugation at 20,000 (RCF) 4°C for 30 minutes.

### Immunoprecipitation

Immunoprecipitations (IPs) were performed on nuclear protein lysates prepared in Nuclear Buffer (prepared as described above). 10μg of antibody was coupled to 50μl packed Protein A beads (Sigma P9424) by incubation in 1ml PBS (0.1% Tween-20) at 4°C rotating overnight. Beads were collected by centrifugation at 1700 × g for 3 minutes and washed twice in 1ml 0.2 m sodium borate pH 9.0. Antibodies were then crosslinked to beads by incubation in 1 ml 0.2 m sodium borate pH 9.0 (20mm dimethyl pimelimidate dihydrochloride) at room temperature rotating for 30 minutes. The reaction was terminated by washing the beads once in 1ml 0. 2M ethanolamine pH 8.0 and incubating for 2 hours at room temperature rotating in 1ml 0.2m ethanolamine pH 8.0. Beads were washed twice in Buffer C100 (20mm HEPES pH 7.6, 0.2mm EDTA, 1.5mm MgCl2, 100mm KCl, 0.5% Nonidet P-40, 20% glycerol) and blocked for 1 hour 4 °C rotating in Buffer C100 with 0.1mg/ml insulin (Sigma, I9278), 0.2mg/ml chicken egg albumin (Sigma A5503), 0.1% (v/v) fish skin gelatin (Sigma G7041). Antibody-crosslinked beads were incubated with nuclear lysates, in the presence of 250U/ml Benzonase nuclease, at 4°C rotating overnight and washed 5 × 5 minutes in Buffer C100 with 0.02% Nonidet P-40. After the final wash, beads destined for immunoblotting were resuspended in 50μl 2× SDS sample buffer. Immunoprecipitated material was eluted by boiling for 5 minutes with shaking, and associated proteins were separated by SDS-PAGE and analyzed by immunoblotting. Beads destined for mass spectrometry analysis were washed once in IP buffer containing 0.02% Nonidet P-40 followed by one wash in IP buffer with no detergent.

### Mass spectrometry analysis

Proteins were treated with trypsin as described (86). Samples were redissolved in 50 μl of Trifluoroacetic acid 0.1% (vol/vol) in water, as buffer A, and sonicated for 1 minute and centrifuged for 15 minutes at 15000 × g. Analysis was carried out on an Ultimate 3000 RSLCnano HPLC system connected to a mass accuracy high resolution mass spectrometry, Q Exactive (ThermoFisher). The MS instrument was controlled by Xcalibur software (ThermoFisher).

Each sample was loaded onto a 75 μm x 15 cm C18 column (particle diameter 1.8 μm, pore size 120 Å) and was separated by an increasing acetonitrile gradient over 100 minutes at a flow rate of 200 nL/min.

MS analysis was done in DDA mode: parent ion spectra (MS1) were measured at resolution 60,000, AGC target 3e6. Tandem mass spectra (MS2; up to 20 scans per duty cycle) were obtained at resolution 15,000, AGC target 2e5 and collision energy of 27.

### Data processing

Data were processed using MaxQuant version MaxQuant version 1.4.3.22 (88) using the human UniProt database (Taxon identifier 9606, Proteome ID UP000005640, Protein Reviewed 20,380). The following search parameters were used: Fixed Mod: carbamidomethylation; Variable Mods: methionine, oxidation; Trypsin/P digest enzyme (maximum 2 missed cleavages); Precursor mass tolerances 6 ppm; Fragment ion mass tolerances 20ppm; Peptide FDR 1%; Protein FDR 1%.

“Label-Free Quantitation; LFQ,” “iBAQ,” and “Match Between Run” settings were selected. Reverse hits and contaminants retrieve from the cRAP database (https://www.thegpm.org/crap/) (89) were filtered out and not considered further.

### Data and Statistical Analysis

Bioinformatic analysis of the MaxQuant output files and data visualization was performed with Perseus software version 1.4 (90) and RStudio employing the following packages: ggplot2 and ggrepel, and clusterProfiler. LFQ values were extracted from the protein group table. No additional normalization steps were performed, as the resulting LFQ intensities are normalized by the MaxLFQ procedure (42).

In Perseus software, the LFQ values were transformed (log2) and a protein was considered quantified only if it was detected at least two out of three biological replicates. Missing values imputation was carried out from a normal distribution (width: 0.3, downshift: 1.8), and a two-tailed t test applied with correction for multiple testing (Benjamini). Volcano plots were constructed using the permutation-based FDR (1%) approach (90) (91), and set the significant differences in the protein abundance (≥1.5-fold change).

Gene ontology analysis was performed using the ‘enrichGO’ function of the clusterProfiler R and Bioconductor package with parameters ‘pAdjustMethod = ‘BH’, ont = ‘BP’, qvalueCutoff = 0.05) (92). Protein alignment was performed using Clustal-Omega with default settings (https://www.ebi.ac.uk/Tools/msa/clustalo/) (93)

### Immunoblotting

Protein lysate was quantified using by the Bradford assay. Subsequently protein lysates were separated on SDS-PAGE gels and transferred to nitrocellulose membranes. Membranes were blocked with 5% non-fat milk or 5% BSA at room temperature for 1 hour and incubated overnight with diluted primary antibody at 4°C. Membranes were then washed and incubated with HRP-conjugated goat-anti-rabbit or mouse IgG secondary antibody for 1 hour at room temperature. Membrane was incubated with enhanced chemiluminescence reagents (Thermo Scientific) followed by exposure to X-ray films. Immunoblotting was performed using the antibodies and conditions listed in Supplementary material XXX.

### Gel filtration column chromatography

The SuperoseTM 6 10/300 GL gel filtration column (GE Healthcare) was equilibrated with one column volume of running buffer (20mM Tris pH 8.0, 10% Glycerol, 175mM NaCl, 0.5mM DTT, 1mM PMSF). 300-500μg of total nuclear protein (prepared as described above) was injected and run through column at 0.35mL/min. 1mL fractions were collected and protein was concentrated by incubation with 4μL StrataClean resin (Agilent Technologies) for 1 hour at room temperature. Resin was collected by centrifugation at 5000rpm for 3 minutes and protein was eluted by boiling in 20μL 2X SDS sample buffer for 5 minutes shaking at 3000 (RCF). Eluted protein analyzed by SDS-PAGE and immunoblotting

### Real-time Quantitative PCR

Extracted RNA was used to generate cDNA by reverse transcriptase PCR using the TaqMan Reverse Transcription kit (Applied Biosytems). Relative mRNA expression levels were determined using the SYBR Green I detection chemistry on LightCycler 480II Real-Time PCR System (Roche). The ribosomal constituent RPO was used as normalizing gene. The primers used are listed in supplemental material.

### Cell Viability Assay

NT2 cells were seeded into the 12-well plate in Dulbecco’s Modified Eagle Medium (DMEM) supplemented with 10% (v/v) Fetal Bovine Serum (Hyclone), 100U/ml penicillin and 100U/ml streptomycin (Gibco).

Lentivirus-infected NT2 were subjected to crystal violet staining using 0.1% crystal violet (CV). After staining, plate wells were subsequently washed with phosphate buffered saline (pH = 7.4) to remove the unbound crystal violet and residual NT2 cells. The plates were then air dried at room temperature and 95% ethanol was added to the wells to resuspend the adhered stained cells. The ethanol bound crystal violet stain of adhered cells were quantified by measured at 590nm in the spectrophotometer.

## Acknowledgements

The research reported was supported by The Comprehensive Molecular Analytical Platform (CMAP) under The SFI Research Infrastructure Programme, reference 18/RI/5702.

## Author Contributions

G.O. Conceptualization, Investigation, Methodology, Visualization, Writing – original draft, Writing – review and editing. K. W and NM. Methodology, Writing – review and editing.

